# Functional and regulatory diversification of *Period* genes responsible for circadian rhythm in vertebrates

**DOI:** 10.1101/2023.03.09.531707

**Authors:** Jun Soung Kwak, M. Ángel León-Tapia, Celian Diblasi, Domniki Manousi, Lars Grønvold, Guro Katrine Sandvik, Marie Saitou

**Author notes:** co-1st authors. Email addresses M. Ángel León < >, Junsoung Kwak < >, Guro Katrine Sandvik < >, Lars Grønvold < >, Domniki Manousi < >, Célian Diblasi < >. **Author Contribution:** Junsoung Kwak: Conceptualization, Data curation, Investigation, Writing – Original Draft Preparation Miguel Ángel León-Tapia: Conceptualization, Data curation, Investigation, Writing – Original Draft Preparation Celian Dilasi: Investigation, Software Domniki Manousi: Software Lars Grønvold: Investigation, Software Guro Katrine Sandvik: Conceptualization Marie Saitou: Conceptualization, Investigation, Writing – Original Draft Preparation Writing – Review & Editing.

## Abstract

The Period genes (*Per*) play essential roles in modulating the molecular circadian clock timing in a broad range of species, which regulates the physiological and cellular through the transcription-translation feedback loop. While the *Period* gene paralogs are widely observed among vertebrates, the evolutionary history and the functional diversification of *Per* genes across vertebrates are not well known. In this study, we comprehensively investigated the evolution of *Per* genes, including de novo binding motif discovery by comparative genomics. We also determined the lineage-specific transcriptome landscape across tissues and developmental stages and phenotypic effects in public RNA-seq data sets of model species. We observed multiple lineage-specific gain and loss events of *Per* genes, though no simple association was observed between ecological factors and *Per* gene numbers in each species. Among salmonid fish species, the *per3* gene has been lost in the majority, whereas those retaining the per3 gene exhibit not a signature of relaxed selective constraint but rather a signature of intensified selection. We also determined the signature of adaptive diversification of the CRY-binding region in *Per1* and *Per3*, which modulates the circadian rhythm. We also discovered putative regulatory sequences, which are lineage-specific, suggesting that these cis-regulatory elements may have evolved rapidly and divergently across different lineages. Collectively, our findings revealed the evolution of *Per* genes and their fine-tuned contribution to the plastic and precise regulation of circadian rhythms in various vertebrate taxa.

**Significance:** The *Period* (*Per*) genes play essential roles in the circadian rhythm in animals. In this study, we comprehensively investigated the evolutionary diversification of the three types of *Period* genes in vertebrates. As a result, we observed a rapid evolution and sub-functionalization of these genes, especially adaptive diversification signatures in the protein-binding region, which plays a crucial role in regulating circadian rhythms. This underscores the fine-tuned contribution of *Per* genes in the biological clock’s precision and adaptability across various vertebrate taxa.

## Introduction

The circadian clock regulates the physiological and cellular timing of organisms, adapting to the external environment with the 24-h light/darkness cycle of the solar day at the molecular level. The system is essential for an organism’s rhythmic behavior and physiological responses, including sleep/wake cycle and foraging. The circadian clock is thus fundamental to the survival and adaptation of organisms in their respective habitats (Cermakian & Sassone-Corsi 2000; Schultz & Kay 2003; Roenneberg & Merrow 2016; Challet 2019). The biological rhythm can efficiently interact with the dynamic and temporal external resources and environments by adjusting its physiological processes (Patke et al. 2020; Umemura & Yagita 2020).

The first genetic factor of circadian rhythm regulation, the *Period* gene, was discovered in *Drosophila* mutant strains that showed unusually long or short biological rhythm patterns (Konopka & Benzer 1971; Bargiello et al. 1984; Zehring et al. 1984). The Period gene has also been reported in other invertebrates, such as insects, German cockroaches (Lin et al. 2002), gastropods (Aplysia and Bulla)(Siwicki et al. 1989), and crustaceans (American lobster) (Grabek & Chabot 2012). Mammalian homologs of the *Period* gene were also characterized soon after that (Tei et al. 1997), and the genetics, molecular mechanism, and behavioral biology of *Period* genes have been extensively studied in model organisms (Wager-Smith & Kay 2000; Cermakian et al. 2001; Pendergast & Friday 2009). These studies revealed that transcription-translation feedback loops regulate the critical role of the molecular circadian clock, core clock genes with changes in external cues, in which *Period* genes (*Per*) play the key role, at least in teleost fish and mammals (Langmesser et al. 2008; Relógio et al. 2011; Idda et al. 2012; Jensen et al. 2012; Lande-Diner et al. 2013; Ono et al. 2017). When there is enough light in the daytime, E-box, the DNA element is bounded by the transcription factors CLOCK-BMAL1, and the level of the PER and Cryptochrome (CRY) proteins are suppressed (Cox & Takahashi 2019). When night comes, and once PER and CRY levels have sufficiently dropped in the dark condition, a new cycle of CLOCK–BMAL1 begins to the *Per* and *Cry* transcription (Vallone et al. 2004; Patke et al. 2020).

Humans and mice have three copies of *PER* genes, *PER1*, *PER2*, and *PER3*, likely due to the two rounds of ancient genome duplications (von Schantz et al. 2006), a prime example of evolutionary processes by ancient gene duplications (Ohno 1970, 1999), highlighting the evolutionary significance of these genes. *Per* paralogues show some sequence similarity and overlapping functions, and specific functions for each gene. In mice, *Per1* and *Per2* genes are responsible for the circadian rhythm, while *Per3* did not indicate an observable, crucial role in circadian rhythm (Bae et al. 2001). On the contrary, a genetic variant in *PER3* in humans is associated with sleep disorders (Zhang et al. 2016). In ray-finned fish, Wang (Wang 2008) highlighted that additional teleost per gene copies originated from a teleost-specific genome duplication, with different species retaining various per duplicates. In that study, the absence of *per3* in sticklebacks (Gasterosteidae) is also reported, indicating a lineage-specific evolutionary pattern of per gene family. Moreover, relaxed selection on *per1b* in Tinaja cavefish was reported (Idda et al. 2012; Mack et al. 2021), while Mack et al. (2021) demonstrated positive selection on per genes in *Squalius* freshwater fish species with population specificity. These studies suggest that the evolution of the *per* gene has diversified in response to ecological niches, contributing to variation in circadian behavior in the biological clock. The rapid turnover of circadian genes in amphibians also indicates the adaptive role of these genes (Stanton et al. 2022). However, despite recent advancements in genomic data, the comprehensive characterization and understanding of the evolutionary history and diversification process of *Per* genes in most non-model vertebrates remain a significant gap in our knowledge.

At the molecular and physiological levels, it has been thought that the *Per* genes are exclusively expressed in the suprachiasmatic nucleus, a brain region in the hypothalamus, which is responsible for controlling circadian rhythms (Takumi et al. 1998; Reppert & Weaver 2002). Contrary to the historical notion, studies revealed that *Per* genes are also expressed in cell lines and non-brain tissues (Balsalobre 2002), sometimes with circadian rhythmic expression patterns, such as in mouse liver (Siepka et al. 2007) cell lines (Ramanathan et al. 2014), and human cell lines (Gabriel et al. 2021). Currently, it is believed that the circadian clock is controlled by the master clock located in the suprachiasmatic nucleus with core neuron gene expressions and is hierarchically conveyed to peripheral clocks in tissues all over the body (Patke et al. 2020; Umemura & Yagita 2020). This revelation highlights the complexity and ubiquity of the circadian clock beyond the central nervous system, indicating a systemic regulation of circadian rhythms. Interestingly, studies suggested that circadian rhythm starts functioning even before birth in various species (Landgraf et al. 2015). For example, differential regulation of *per2* and *per3* expression during zebrafish embryogenesis is reported (Delaunay et al. 2003). It is reasonable that circadian genes are expressed in oviparous embryos with a transparent eggshell, which is exposed to light in the day and darkness in the night cycles. Studies indicated that circadian rhythm genes are expressed even in mammal embryos that cannot directly sense the day/night cycle by light (Shimomura et al. 2001; Yagita et al. 2010), and it is speculated that the mother transmits the fetus information about the day/night cycle so that the fetus can calibrate the circadian rhythms (Seron-Ferre et al. 2007; Landgraf et al. 2015). In addition, *Per* genes have non-circadian functions, such as DNA damage response (Fu et al. 2002) and embryonic neuron development (Nagata et al. 2019) underscores their multifaceted roles in biological processes. Despite numerous studies on *Per* expression in various conditions, the simultaneous investigation of *Per* gene paralogs remains unexplored. Thus, the understanding of their shared and species-specific roles is still yet to be investigated. This paper aims to bridge this gap by comprehensively investigating the evolutionary history and the functional diversification of *Per* genes in vertebrates.

## Results and Discussion

### Lineage-specific gene gain and loss of Period genes in vertebrates

First, we curated the *Per* genes in vertebrates (**Fig 1A**) using Ensembl Compara v106 and then visualized the evolutionary history of *Per* gene gain and loss in the species-based phylogenetic tree. After genome quality filterings (**Materials and Methods**), we included 154 species in total and obtained the sequences of 173 *Per1* genes, 212 *Per2* genes, and 150 *Per3* genes. There is no correlation observed between the reported *Per* gene number and the genome quality metrics (N50, **Supplementary Table1**) (rho= - 0.00038, p > 0.05, Spearman correlation), excluding the possibility that the genome quality could influence the observed patterns of *Per* gene numbers. The phylogenetic signal (Pagel’s lambda λ) (Pagel 1997) was high in the three *Per* genes reflecting a pattern where closely related species tend to be more similar in *Per* number than the more distantly related species. *Per1* and *Per 3* showed the strongest phylogenetic signal, and *Per2* also showed a high phylogenetic signal: *Per1* λ= 1.00, *Per2* λ= 0.94, and *Per3* λ= 1.00. Most mammalian and amphibian species have three paralogs (*Per1-3*) (**Fig 1A**), one for each family, yet we observed several lineages with marked gene gain/loss. Most reptiles lost the *Per1* gene, while birds lost the *Per3* gene, respectively **(Fig 1B**). The *Per2* number is the most stable among the three genes across taxa, and most species maintain the *Per2* gene, suggesting that *Per2* plays a major, universal role. The consistent presence of the *Per2* gene in the genomes across different taxa underscores its indispensable and universal significance. Cartilaginous fishes have been inferred to be basal extant jawed vertebrates, and their subclass *Elasmobranchii*, including Elephant shark (*Callorhinchus milii*), possess only one *per2* gene. In contrast, in most teleost fish species, two *per2* genes were observed, likely due to the teleost fish-specific whole genome duplication event (Ts3R) (Near et al. 2012; Berthelot et al. 2014; Inoue et al. 2015). In addition, there was a *Cyprinidae*-specific whole genome duplication (Cs4R) (Xu et al. 2019) and a *Salmonidae*-specific whole genome duplication (Ss4R) (Lien et al. 2016), which led to additional duplications of *Per* genes in each lineage **(Fig 1C**). Conversely, *per3* gene loss is observed in multiple distant fish lineages, including pike (*Esox lucius*), most salmonid species (Bolton et al. 2021), and cod (*Gadus morhua*), (**Fig 1C**). Although the loss of another circadian gene and retrogression of circadian rhythm is reported in cavefish (*Astyanax mexicanus*) (Mack et al. 2021), there is no specific trend observed in the *per* gene copy number in cavefish. We observed that *per3* genes are lost in most salmonid species, except for Atlantic salmon *(Salmo salar)*, and brown trout *(Salmo trutta)*, consistent with a previous study (Bolton et al. 2021). To test if *per3* gene is under relaxed selection in these two species, we used a codon-based statistical test to detect relaxed selection, RELAX (Wertheim et al. 2015), using the *per3* genes in these three species with retained *per3*, compared to the Acanthomorphata group. As a result, we observed a signature for intensified positive selection in the *per3* genes in these species with retained *per3*, compared to the control group (k=1.28, p=0.023) (**Fig S1**). This implies that the *per3* gene maintains indispensable roles in salmonid species that have this gene, which is contrary to the current view that *Per is the least important, based on earlier knock-out mouse studies, and thus, least studied among the paralogs* (Bae et al. 2001).

**Fig 1.**
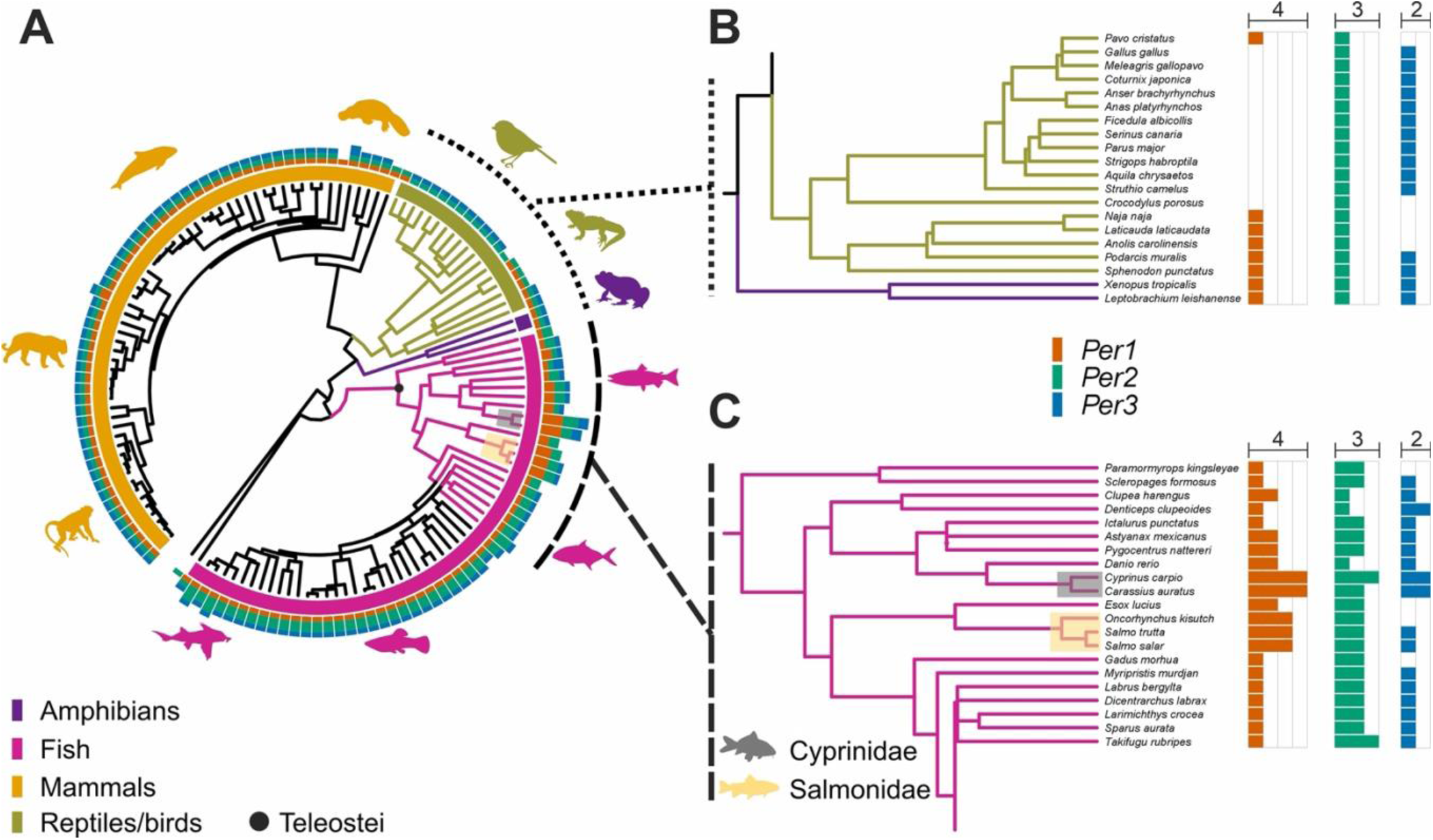
The evolutionary history of the *Period* gene (*Per*) gain and loss in vertebrates. **A**. Phylogenetic tree and numbers of *Per* paralogs in vertebrates. Bar colors around the tree denote the main vertebrate taxonomic clades; amphibians in purple, fishes in deep pink, mammals in gold, and reptiles/birds in olive green. The bar color after the tips indicates the gene number for each species; orange for *Per1*, dark green for *Per2*, and blue for *Per3*. **B**. Zoom in on amphibians, reptiles, and bird clades showing unusual gene loss patterns. **C**. Zoom in on *Salmonidae, Cyprinidae, and Chichilidae* clades that show unusual gene gain-loss patterns.

We also reconstructed ancestral *Per* states calculating for each gene states transitions based on an equal rate (ER), all-rates-different (AR), and symmetrical (SYM) using the hidden-rate model (Beaulieu et al. 2013). The best model of ancestral state transitions for *Per1* was ER (logL = -47.46), ARD for *Per2* (logL = -34.10), and SYM for *Per3* (logL = -45.26). Ancestral state estimations showed a similar pattern for the three *Per* genes in the vertebrate’s phylogeny, in which the current observed gene number was derived from a single ancestral copy (**Fig 2**). Dynamic gain/loss pattern changes were more recent clades in *Per1* **(Fig 2A**) and *Per3* (**Fig 2C**) than the duplication pattern noticed in *Per2* (**Fig 2B**). Disparity Through Time (DTT) (Harmon et al. 2003) plots for each *Per* gene revealed differences in the patterns of evolutionary history (**Fig 2D**). *Per1* showed a major increase in the among-clade disparity while *Per2* showed stability, likely driven mainly by the duplication in the whole fish clade, with some slight peaks of within-clade disparity led by the diversification in Cyprinidae, indicating that diversification was high later in the evolutionary history of the group. *Per3* had a pattern of gradual increase in the among-clade disparity.

**Fig 2.**
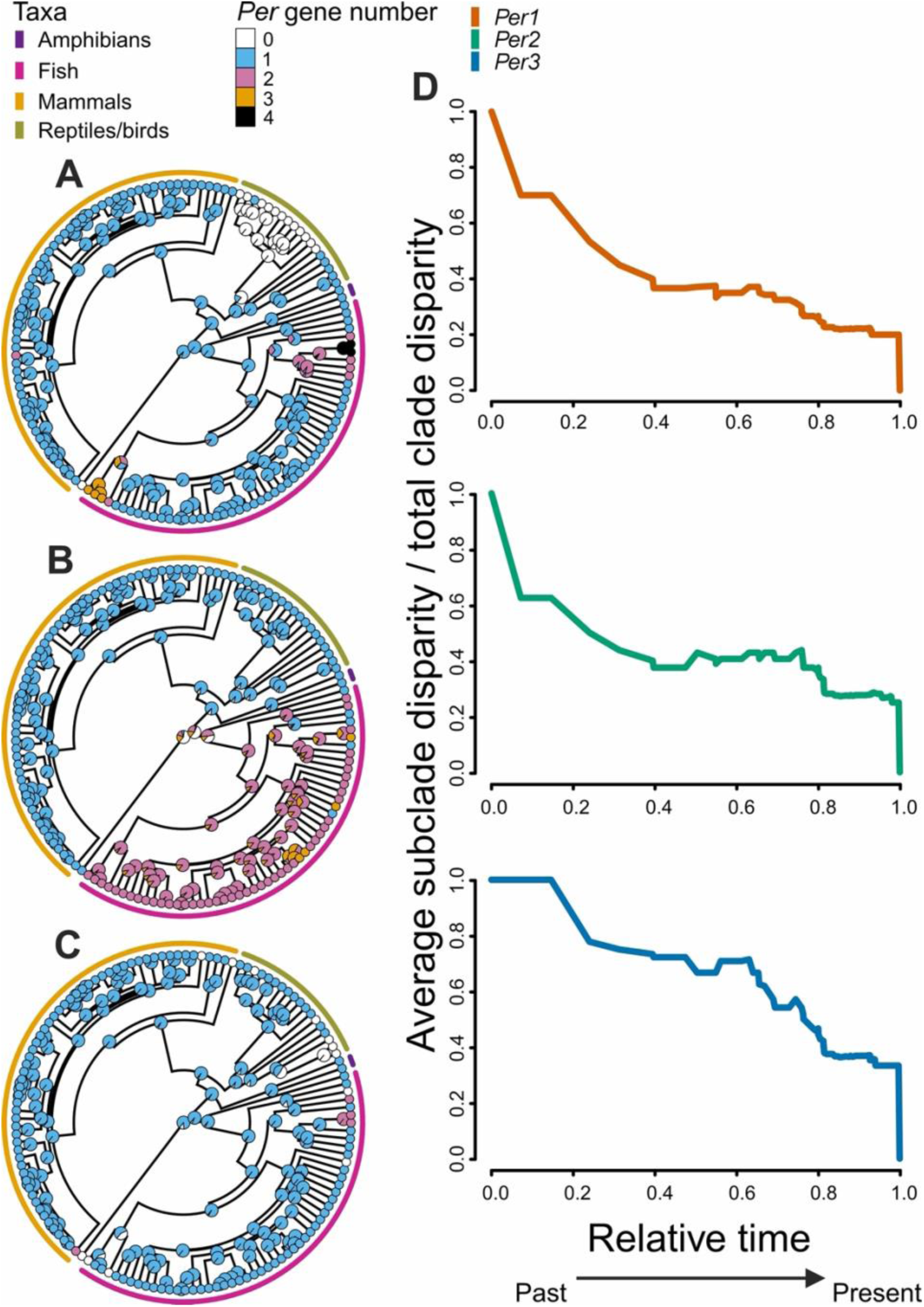
Ancestral state reconstruction for Period genes (*Per*) gain/loss numbers in vertebrates, **A**. *Per1*, **B**. *Per2*, and **C** *Per3*. **D**. Disparity Through Time (DTT) plots for each *Per* gene. The horizontal axis represents relative time values, 0.0 represents the root, and 1.0 the tips of the tree. The vertical axis represents the average subclade disparity divided by the total clade disparity and is calculated by each internal node of the tree. Higher values of disparity correspond to greater variance values within subclades relative to the disparity of the whole subclade.

The evolution of circadian rhythm has been discussed in the context of ecological adaptation and behavioral strategies in both terrestrial and aquatic species, including temperature, diet, habitat, reproduction, and migration; although the effect of these environmental and ecological factors on each species is complex, mutually influential, and variable (Horton 2001; Bloch et al. 2013; Oliveira et al. 2016; Hewitt & Shaikh 2021). We hypothesized that some environmental factors may have driven the evolution of *Per* gene gain and loss (**Material and Methods**). To investigate the evolutionary association between each *Per* gene number and in each species, we conducted the Phylogenetic Generalized Least Squares test (Symonds & Blomberg 2014) between per gene copy number and environmental/ecological factors of 138 species with data available. A total of ten environmental/ecological factors we tested included elevation, temperature, precipitation seasonality, solar radiation, windspeed, water vapor pressure, and net primary productivity, and ecological values included migration, body mass, and habitat ( **Materials and Methods**). To avoid noise due to taxa-specific factors, we separated each taxon (mammals, birds and reptiles, amphibians, and fish) in the analysis (**Materials and Methods**). As a result, we did not observe any statistically significant association between *Per* gene numbers and each environmental or ecological variable (Phylogenetic Generalized Least Squares test, *p-*value less than 0.05 was considered to be significant). We assume that influence of environmental and ecological factors on circadian rhythm is based on a more intricate system. For instance, as a result of a spatiotemporal regulation of the enzymatic activity, rather than the simple *Per* gene copy number. In mammals, compelling evidence underscores the presence of circadian synchrony between predator-prey activity patterns (Caravaggi et al. 2018). These intricate interspecific relationships have emerged as pivotal factors, propelling organisms towards physiological and behavioral adaptations (Bennie et al. 2014). These adaptations, in turn, may wield significant influence as selective pressures in vertebrates, instigating genetic changes within species and shaping their capacity to exploit available resources effectively.

We also examined if the copy gain and loss pattern of each *Per* gene family deviated from other gene families available at Ensembl v104 compara gene trees, as previously reported (Gundappa et al. 2022). We calculated the coefficient of variation of gene numbers of each gene, and found three gene groups, with (A) low variation, (B) medium variation, and (C) high variation. *Per* genes fell in the medium variation cluster. Genes with very low variation included highly conserved essential genes such as Kinesin family-like genes and genes with high variation included dynamically evolving genes, including olfactory receptors (**Fig S2, Table S5**).

### Evolutionary conservation and divergence in *Period* genes function across taxa

Previous studies with intensive genetic engineering and behavioral experiments, primarily in mice, suggested that the *Per3* gene has no intrinsic effect on circadian rhythms compared to the *Per1* and *Per2* genes (Bae et al. 2001), and *Per3* is often excluded from later investigations and discussions probably for this reason (Chiou et al. 2016; Cox & Takahashi 2019). However, this makes it difficult to reveal the comprehensive dynamics of how *Per* gene paralogs share roles, and thus how conserved these roles are across species is not known. To investigate if each *Per* gene function is shared or specific and how much they are evolutionarily conserved, we curated and re-analyzed published phenotype datasets. We first inferred that vertebrate *Per* genes are likely to have a single ancestor by constructing a phylogenetic tree based on the amino acid sequences of Per from vertebrate species, with the *Drosophila* Per gene sequence. The *Drosophila* Per sequence was clustered with the Per3 group, suggesting that *Per3* is the most ancestral form of the *Per* gene family (**Fig 3A**). It is speculated that the ancestral *Per3* gene was duplicated twice due to the first and second vertebrate whole genome duplications and further duplicated by the teleost-specific whole genome duplication corresponding with previous studies (Wang 2008). The phylogenetic analysis also supported that lineage-specific whole genome duplications in common carp (*Cyprinus carpio*) and salmon (*Salmo salar*) have also likely contributed to their additional *per* gene duplications.

**Fig 3.**
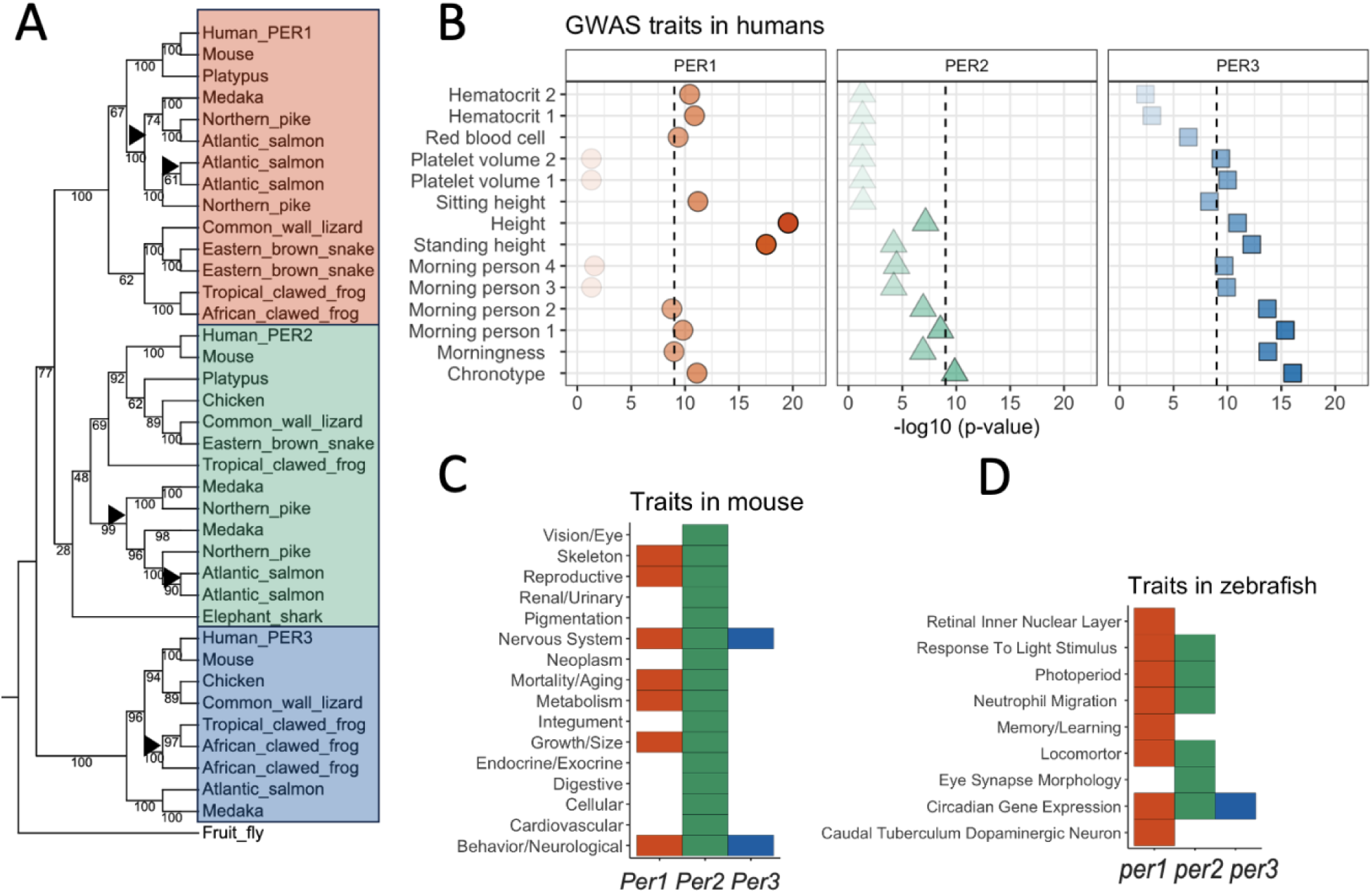
Phenotypic effect differentiation of *Period* genes (*Per*) in various vertebrate species. **A**. Amino-acid-based phylogenetic tree of vertebrates with the outlier fruit fly (*Drosophila melanogaster*) *per*. Amino acid alignments were performed using MAFFT v7 (Katoh et al. 2002). For the tree construction, the neighbor-joining method was applied (Saitou & Nei 1987), focusing on conserved sites across 363 amino acids. The nucleotide substitution model employed was the Jones-Taylor-Thornton model (Jones et al. 1992). Bootstrap analysis was conducted to assess the robustness of the tree, with the number of resampling events set to 100. Bootstrap value is shown in the tree (**file S3**). Black triangles indicate whole genome duplication events. The background color shows the gene classification (Red: *Per1*, Green: *Per2*, Blue: *Per3*). **B.** Reported phenotypes associated with *PER1*, *PER2*, and *PER3* in humans at GWAS Atlas (Watanabe et al. 2019). The -log_10_ p-value of the association between genes and phenotypes is plotted on the x-axis. Only phenotypes with -log_10_ p > 9 in at least one of the *PER* genes are shown. **C**. Reported traits in *Per* deficient mouse (*Mus musculus*) from Mouse Genome Database (MGD), 2021 (Blake et al. 2020). **D.** Reported traits in *per* deficient zebrafish (*Danio rerio*), the Zebrafish Information Network (zfin.org), 2022 (Bradford et al. 2022).

We next investigated the regulatory and functional differentiation between *Per* paralogs by analyzing public gene-phenotype data sets of multiple species. We examined the genome-wide association study hits of variants in the three *PER* genes in humans that are curated and summarized at GWASATLAS (Watanabe et al. 2019). The gene-level p-value of association with traits was computed by MAGMA (de Leeuw et al. 2015), a computational method based on a multiple linear principal components regression model to aggregate the effects of variants on traits in a gene. We extracted traits that showed significant association (*p*-value smaller than 10^-9^) in any of *PER* genes in the human GWAS database (**Fig 3B**). Human *PER1* showed the highest statistical significance with height and medium significance with chronotype and blood-related traits, such as hematocrit, and the percentage of red cells in the blood. *PER2* is also moderately associated with chronotype. *PER3* showed the highest statistical significance with chronotype and also showed some association with height and another blood-related trait, platelet volume. A deletion (54 bases) polymorphism was reported in the genomAD paper (Collins et al. 2020), which is associated with sleep (Archer et al. 2018). This variant is not reported in the 1000 Genomes phase 3 dataset (The 1000 Genomes Project Consortium et al. 2015) and the updated 1000 genome dataset (Byrska-Bishop et al. 2022) as it falls in a complex repetitive region, and little is known about the molecular function and evolutionary significance of the repeats.

It is reported that single *Per1* or *Per2* knockout mice have shorter circadian periods with reduced precision and stability, and both *Per1* and *Per2* knockout mice are completely arrhythmic in constant conditions (Cermakian et al. 2001; Zheng et al. 2001; Bae et al. 2001). Interestingly, *Per3* knockout mice have relatively moderate changes in circadian rhythm, observed light-dependent phenotypes, and reduced body size, growth, and metabolism (Archer et al. 2018; Pereira et al. 2014). In our meta-analysis, using data from Mouse Genome Database (MGD), 2021 (Blake et al. 2020), *Per2* gene has a broad effect on various phenotypes, while *Per1* and *Per3* are only involved in several neurological traits (**Fig 3C**). In zebrafish (*Danio rerio*), based on the Zebrafish Information Network (zfin.org), 2022 (Bradford et al. 2022), it is reported that *per1b* or *per2* deficiency has effects on locomotor, photoperiod, neural development, and neutrophil migration, a blood-related trait similar to the human case, interestingly. The *per1b* gene has specifically reported phenotypic effects such as memory, and *per2* has specifically reported phenotypic effects such as eye synapse morphology. Meanwhile, there are no marked effects reported for *per1a* and *per3*, except for some expression changes of circadian-related genes (**Fig 3D**). These observations indicate the flexibility of each *Per* gene’s roles between species.

### Dynamic evolution of *Period* gene expression pattern between paralogs and species

To understand the regulatory differentiation that may underline such diversity in phenotypic effects of *Per* genes across species, we searched for the *Per* gene expression in various tissues in vertebrates. We compared adult *PER* gene expressions of humans (*Homo sapiens*), from the GTEx portal (GTEx Consortium et al. 2017), mouse (*Mus musculus*) expression from the mouse Gene Expression Database (GXD) (Baldarelli et al. 2021), and African clawed frog (*Xenopus laevis)* Xenbase (Fortriede et al. 2020) **(Fig 4A-C).** In humans, *PER1* showed the highest expressions among *PER* genes overall except for Cerebellum. Notably, *PER3* shows a marked expression level at the Cerebellum among other organs, which is strongly influenced by the sleep-wake cycle (Canto et al. 2017) while it does not show such high expression in other parts of the brain. The *Per2* gene shows mostly low expressions in both the human and the clawed frog, but in the mouse, the *Per2* gene shows a moderate expression level and its expression pattern across species is markedly similar to the *Per3* gene expression. Two *Per3* gene duplicates in the clawed frog, which experienced allotetraploidization around 17–18 Mya (Session et al. 2016), showed similar expression patterns each other across tissues **(Fig 4C)**. The orthologous genes, such as *PER2* in humans and *Per2* in mice, did not necessarily show a strong correlation in their expression pattern between species (r=-0.07, Spearman’s correlation, **Fig 4D)**, suggesting the dynamic evolutionary plasticity of the expression pattern. Rapid transition of tissue-specificity of gene expression has been reported (Fukushima & Pollock 2020), in particular between the ovary and testis, as we also observed in *PER1* in the human and *Per1* in the mouse **(Fig 4AB).**

**Fig 4.**
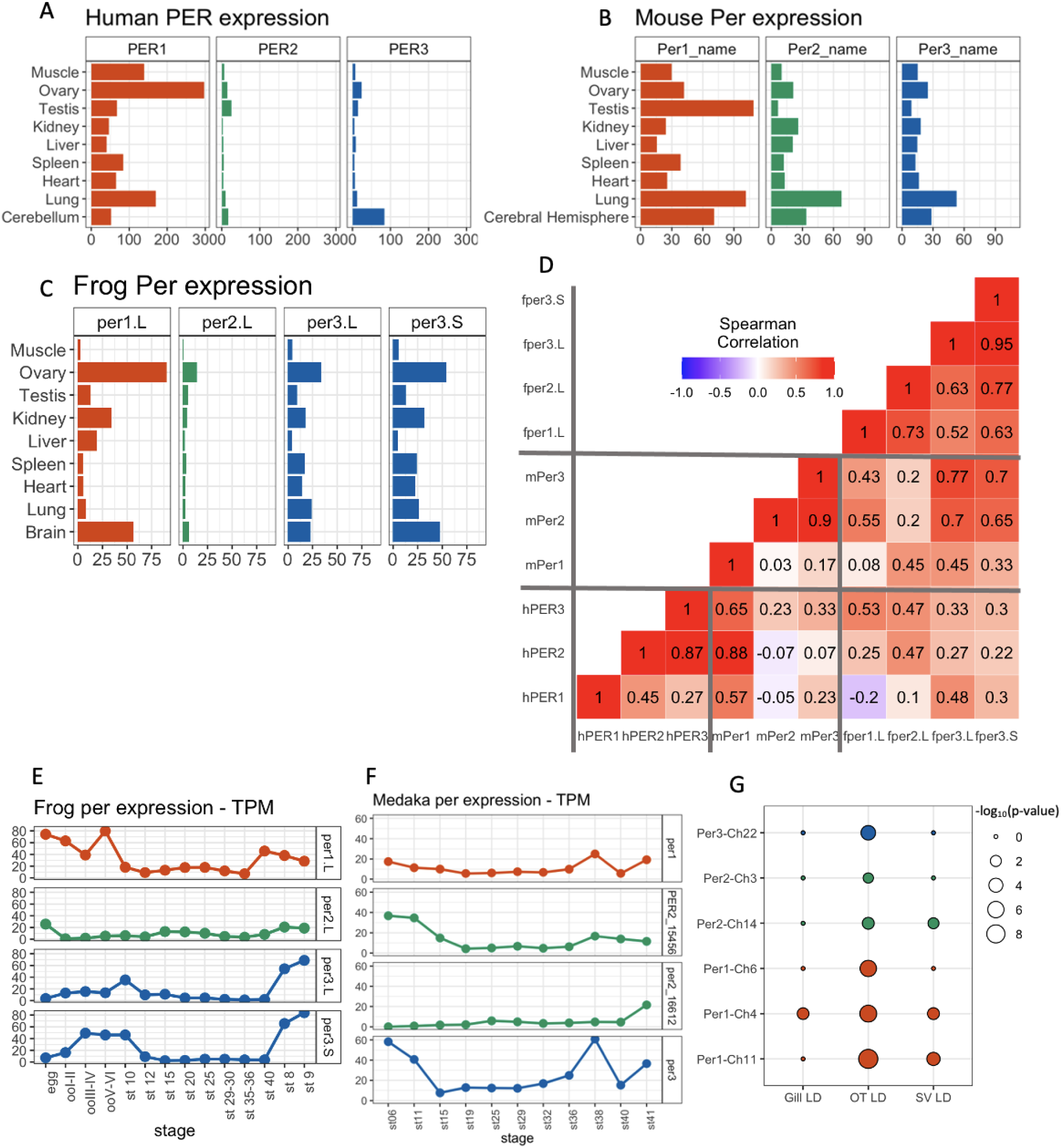
Transcriptomic differentiation of *Period* genes (*Per*) in **A**. Human, **B**, Mouse (*Mus musculus*), and **C** Frog (*Xenopus laevis*). The expression data was obtained from the Genotype-Tissue Expression (GTEx) Portal, the Mouse Genome Database, and the Zebrafish Information Network. **D** Correlation of gene expression patterns between per genes (hPER: human, mPer: mouse, fper: frog). **E**. Developmental change of *per* gene expression in African clawed frog (*Xenopus laevis)* Xenbase (Fortriede et al. 2020). **F**. Developmental change of *per* gene expression in medaka (*Oryzias latipes*), at Medaka Omics Data Portal (Li et al. 2020) **G**. Rhythmicity of *per* genes in tissues in Atlantic salmon (*Salmon salar*), *Optic tectum*, gill, and *saccus vasculosus*. Data is retrieved from supplementary table 8 of (West et al. 2020), one gene with no expression is not displayed.

To further investigate the evolutionary trajectory of this gene expression regulation, we compared the expression pattern of relatively new *Per3* gene duplicates in the African clawed frog, *Xenopus laevis*, which experienced allotetraploidization around 17–18 Mya (Session et al. 2016) and the Japanese medaka (*Oryzias latipes*) which has likely duplicated the *per2* gene because of the teleost genome duplication around 310 Mya (Davesne et al. 2021). The newly duplicated two *Per3* genes in *Xenopus laevis*, showed a very similar expression pattern both across tissues and across developmental stages Data from Xenbase (Fortriede et al. 2020) (**Fig 4E**), such as high expression in the ovary, liver, brain, and lung, and stage 8 and 9, showing their regulatory mechanism is still very similarly conserved. On the contrary, the two Japanese medaka *per2* genes were already differentiated at the regulation level at Medaka Omics Data Portal (Li et al. 2020) (**Fig 4F**). Collectively, the *Per* gene expression pattern in tissues is not necessarily conserved across species and is dynamically diversifying, probably due to the rapid evolution of gene regulation (Rifkin et al. 2005; Hill et al. 2021). We further asked about the developmental pattern of *PER* gene in different parts of the human brain using the Brainspan dataset (https://www.brainspan.org/) (Johnson et al. 2009; Kang et al. 2011) **(Fig S3)**. Overall, *PER1* showed the highest expression across tissues, especially in the 37 post-conception weeks fetus, except for the cerebellar cortex. *PER3* shows high expression in the cerebellar cortex after birth when humans are exposed to the external light and dark cycle. *PER3* also showed high expression in the thalamus, which acts as a sensory hub (Hwang et al. 2017). Neither the suprachiasmatic nucleus nor the broader region, the hypothalamus, which is responsible for controlling circadian rhythms (Takumi et al. 1998; Reppert & Weaver 2002), are reported in Brainspan. In considering the function of *Period* genes, we need to consider their circadian rhythms. Although there have not been many reports on the rhythmicity of all the *Per* genes in one study, West et al. (West et al. 2020) measured the expression of circadian gene paralogues in Atlantic salmon during development in various tissues by using JTK cycle. We extracted the rhythmicity of *Period* gene expression from West et al. (West et al. 2020) (**Fig 4G**). In the Optic tectum, all the expressed *per* genes showed rhythmicity, while in saccus vasculosus, three of seven salmon *Period* genes showed rhythmicity. In gill, only *per1* on chr4 showed moderate rhythmicity.

### Evolutionary conservation and diversification at the sequence level

*Per1*, *Per2*, and *Per3* do not necessarily show the same phenotype and expression pattern between species, rather, these genes appear to be evolving their expression patterns rapidly. To investigate the genomic basis of this dynamic evolution of the *Per* gene regulation, we investigated how the coding sequences and putative regulatory regions of *Per* genes evolved.

We first explored if there are any sites under diversifying selection on *Per1*, *Per2*, and *Per3* vertebrate genes independently, with MEME (Mixed Effects Model of Evolution) (Murrell et al. 2012) on Datamonkey v2.0 (Weaver et al. 2018) (**Table S6**). MEME estimates a site-wise synonymous and a two-category mixture of non-synonymous rates and uses a likelihood ratio test to determine the signal from episodic diversification, a combination of strength of selection and the proportion of the lineages affected. As a result, we observed signatures of diversifying selection in the CRY-binding sites in *Per1* (site “2566” in **Table S6** for all species and site “1583” in **Fig 5B** with the selected species) and *Per3* (site “2102” in **Table S6** for all species and site “1393” in **Fig 5B** with the selected species), as shown in **Fig 5AB** and **Fig S4,** contrary to the general notion that protein binding sites in a gene are evolutionarily conserved (Guharoy & Chakrabarti 2010). As summarized in the introduction section, the PER and Cryptochrome CRY proteins orchestrate the circadian feedback loop with the CLOCK–BMAL1 (Vallone et al. 2004; Patke et al. 2020). CRY-binding region of the *Per* gene was characterized in rats (*Rattus norvegicus*) (Miyazaki et al. 2001). The site under selection in CRY-binding region is conserved, but after that, the latter binding region shows a dramatic decline in conservation (**Fig 5A and 5B)**, which may indicate sub-functionalization between paralogs, and the molecular basis of functional diversification of *Per* genes between species. *Per2* gene showed a marked signature of diversifying selection around the site 263 in the alignment (**Fig S4, TableS6),** while we did not find known functional annotation there but “Polar residues, Compositional bias” in Uniprot (corresponding with the 46th “N” in NE**<u>N</u>**CS in the human sequence, O15055 · PER2_HUMAN at UNIProt, https://www.uniprot.org/uniprotkb/O15055/entry, last accessed, July 3^rd^ 2023 (Deutsch et al. 2023).

**Fig 5.**
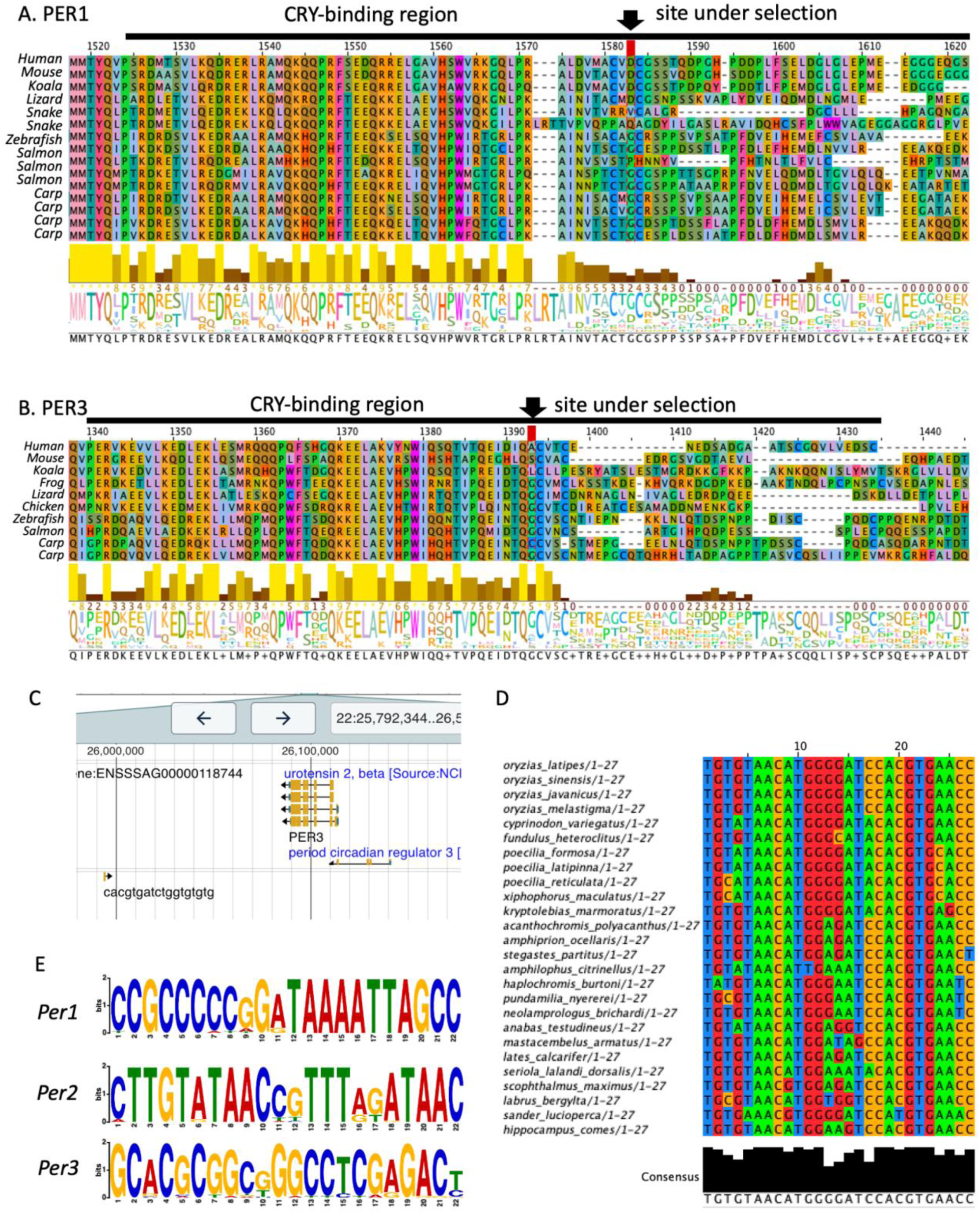
**AB**. The site with diversifying selection detected (arrow) by MEME (Mixed Effects Model of Evolution) (Murrell et al. 2012) on Datamonkey v2.0 (Weaver et al. 2018), which are both located in the CRY_binding sites of *Per1* (**A**) and *Per3* (**B**). As a result, we observed signatures of diversifying selection in the CRY-binding sites in *Per1* (site “1583” in **Fig 5A**) and *Per3* (site “1393” in **Fig 5B** with the selected species). The sites in Fig 5 and **Table S5** do not have the same numbers because only selected species are aligned and shown in Fig 5 for a simplified visualization purpose, and **Table S5** is based on the aligned sequences of all the investigated species. **C**. A sequence, which is similar to the mammalian *PER1*-regulatory sequence, CACGTGatctggTGTGTG, near the *per3* gene in Atlantic salmon (*Salmo salar*) on Salmobase, (Ssal_v3.1, 22:25993822-25993839). **D**. Highly conserved *per2* regulatory region of Japanese medaka (*Oryzias latipes*) HdrR (ASM223467v1) 5:28606354-28606380 region in *Percomorpha.* **E.** MEME (Multiple Expectation maximizations for Motif Elicitation) *de novo* motif discovery results using the 1000 base upstream sequences of each *Per* gene on MEME Suites (Bailey et al. 2015). Putative motif sequence with the top bits score for each gene is displayed. *Per1*: CCGCCCCCGGATAAAATTAGCC (found in *Eutheria*), *Per2*: CTTGTATAACCGTTTAGATAAC (found in *Teleostei*), *Per3*: GGARGCCTAAATATAGGAGGCG (found in *Boreoeutheria*)

We also observed variations in gene lengths, which are presented in **Table S7**. While a subset of genes appeared short, establishing a definitive cutoff for functional loss based solely on gene length proved challenging. Consequently, we expanded our investigation to include domain analyses to identify the presence of conserved functional domains as indicators of potential gene activity. Interestingly, some genes, such as the per3 gene in sheep (ENSOARG00020020217) without recognizable domains reported (**Table S8**), still showed rhythmic expression patterns, suggesting functionality (Varcoe et al. 2014). This underscores the complexity of inferring gene pseudogenization from domain absence or gene length alone.

We further investigated the conservation and putative regulatory regions of *Per* genes. There have been multiple studies that conducted a comparative analysis of the *Per* gene regulatory region. Nakahata et al. (Nakahata et al. 2008) proved that two tandem E-box-like sequences in tandem with a six-base-pair interval are necessary for cell-autonomous oscillation in the mammal and are conserved in various mammalian species. Here, we explored the conservation of E-box elements, expanding their investigations. At the same time, Paquet et al. (Paquet et al. 2008) proposed a model to predict the CLOCK-controlled cis-elements with E-boxes that regulate *Per* gene expression in various species, from *Drosophila* to mammals; Paquet et al (Paquet et al. 2008). also indicated that the E-box sequence is highly conserved between fish and mammals.

As general Blast search is not fine-tuned for searching such motif sequences with short query sequences with very rapid turnovers, we used Jbrowse (Buels et al. 2016; Diesh et al. 2022) -based SalmoBase (Samy et al. 2017) (https://salmobase.org/, last accessed, January 5^th^ 2023) to investigate the promoter sequences, allowing ambiguous spacer sequences around the *per* genes in Atlantic salmon. We searched for the three mammalian sequences reported in Nakahata et al. (Nakahata et al. 2008), allowing the six-base spacer sequences for against 100K upstream and downstream within each *Per* gene region. Of the combination of three query sequences and seven salmon *per* genes, we only found one *per1-regulatory-like* sequence, CACGTGatctggTGTGTG, near the *per3*, but over 80K apart from the gene, and in a repeat region (**Fig 5C**). In Ensembl Comparative Region Comparison Analysis using the 65 fish EPO-Extended, the sequence (Ssal_v3.1, 22:25993822-25993839) was observed only in the Atlantic salmon genome and Brown trout (*Salmo trutta*) genome. It is thus plausible that the mammalian *Per1*-regularoty-like sequence has arisen independently from mammals in this lineage, and the ancestral sequences underwent rapid turnover and are not detectable. The sequence that Paquet et al. (Paquet et al. 2008) estimated corresponds to Japanese medaka HdrR (ASM223467v1) 5:28606354-28606380 region, near the *per3* gene, while it is located near the *Per2* sequence in mammalian species (Paquet et al. 2008). In Ensembl 65 fish EPO-Extended, this region is widely observed in *Percomorpha* (**Fig 5D**) as represented in Paquet et al. (Paquet et al. 2008). However, the zebrafish (GRCz11, 2: 48,352,711-48,352,738) sequence is located near the *per2* gene, and this region was not detectable in other species. This implies that at least the zebrafish sequence evolved to become similar to the regulatory sequence in its lineage, independent from other lineages. These observations of salmon and zebrafish sequences resemble *per* regulatory regions, suggesting the fast rise and fall of the regulatory sequences concordant with previous observations on the evolution of regulatory elements in general (Stone & Wray 2001; Neme & Tautz 2016).

To uncover potential cis-regulatory elements associated with the per1, per2, and per3 genes, we conducted a comprehensive, unbiased *de novo* motif search for 1000 base upstream sequences of per1, per2, and per3 genes using the MEME suite (Bailey & Elkan 1994; Bailey et al. 2009, 2015). Our analysis aimed to identify enriched sequences that could potentially serve as cis-regulatory elements. While MEME identified 10 sequences with the lowest *p-*values, we also observed that some of the sequences were merely repetitive sequences, which are likely false positives and unlikely to be cis-regulatory mechanisms. Therefore, we chose to focus on the sequences with the highest bit scores per gene. The putative regulatory sequences we identified are as follows (**Figure 5E, Table S9**). *Per1*: CCGCCCCCGGATAAAATTAGCC (found in *Eutheria*), *Per2*: CTTGTATAACCGTTTAGATAAC (found in *Teleostei*), *Per3*: GGARGCCTAAATATAGGAGGCG (found in *Boreoeutheria*). These results highlight the lineage specificity of these sequences, suggesting that these cis-regulatory elements may have evolved rapidly and divergently across different lineages. Collectively, the rapid sequence and regulatory evolution that we observed, despite the relatively stable gene copy number, are likely to contribute to the dynamic effect of the *Per* gene family on the phenotype, both circadian and noncircadian effects in various species.

## Conclusion

One of the major questions in evolutionary genomics is how duplicated genes develop the diverse phenotypes in various species by fine-tuning their original functions. This study concentrated on the evolutionary trajectory and functional diversification of the *Per* gene family in vertebrates, key regulators of circadian rhythms. We unveiled how duplicated copies of *Per* genes have adapted their functions across species. While *Per1*, *Per2*, and *Per3* paralogous genes were duplicated in the common ancestor of vertebrates. Notably, most birds have lost the *Per1* gene, whereas the majority of reptiles have shed the *Per3* gene. The *Per2* gene, however, has been retained across most species, suggesting its crucial, universal role in circadian regulation. Contradicting the prevailing perception that *Per3* is of lesser importance (Bae et al. 2001), our analysis indicates intensified selection of *Per3* in certain salmonids, as opposed to relaxed selection. This challenges the current understanding and underscores the gene’s significant yet overlooked role. Through comparative analysis of public genome, phenotype, and transcriptome datasets, we determined that the phenotypic impacts and expression patterns of *Per* gene paralogs exhibit variability across species and paralogs. This variability underscores the rapid evolution of sub-functionalization within the *Per* gene family, indicating a dynamic evolutionary response to diverse environmental and physiological demands.

We postulate that as more transcriptome and gene-phenotype datasets of a variety of species become available, not only genome datasets but the evolution of gene function and regulation will be more comprehensive in the context of how genome evolution contributes to cross-species phenotypic diversity through molecular mechanisms. Although knockout/knock-in studies are a powerful way to understand gene function in a controlled environment, the effects of each *Per* gene are often not investigated within one study. Therefore, when we encounter a lack of known phenotype report of a gene compared to another, it is difficult to know if this is because that gene was “not investigated” or “was examined, but no observable changes were found/thus not reported”, which remains a limitation to investigating the cross-species effects of the paralogs. Our study represents the first step in integrating multiple angles of -omics datasets to understand the functional effect of each *Per* gene in multiple species. It is important to note that although there are interesting lineage-specific gene loss trends in reptiles and birds at the genomic level, currently, there are not many comprehensive cross-tissue transcriptome data and gene-phenotype data available in these taxa. Also, in most transcriptomic studies, the sampling time is not considered and reported, which may raise limitations for comparative circadian rhythm gene investigation. While large-scale international genome sequencing projects for various non-model organisms are underway, the next step for evolutionary genomics is to connect insights from genome information to their functional roles on molecular and macroscopic phenotypes and environmental conditions.

## Material and Methods

### Gene Sequences

We have retrieved all available *Per1*, *Per2*, and *Per3* sequences from Ensembl Compara v106, via the Wasabi app (http://wasabiapp.org/). We initially obtained 200 species. To filter out species with potential low-quality genomes, we removed 18 species with sequences containing “N” (script available in Supplementary Material). We also excluded 25 species that have any *Per* genes not starting from methionine code, which suggests patchy gene annotations. We further removed 10 species with inconsistent reports, which had zero *Per1*/*Per2*/*Per3* genes in Ensembl Compara but with *Per1*/*Per2*/*Per3* existing in NCBI orthologs (last accessed, August 15t^h^ 2022) and two species with projection-built genomes. We tested Spearman’s correlation between the observed Per gene number and the genome quality metrics (N50, obtained from NCBI). We examined the copy gain and loss pattern of *Per* gene in relation to the other gene families, calculating the coefficient of variation for each one using cmstatr package (https://doi.org/10.21105/joss.02265) in R software v4.2.2 (https://www.R-project.org/).

To inspect likely pseudogenes, we removed branches of gene trees that were significantly longer than other genes (likely because they accumulated mutations at a faster rate than others, which is a sign of pseudogenization) by using TreeShrink (Mai & Mirarab 2018), a software that detects branches that significantly increase the tree diameter. We set the chance of false positives to 10 percent. We thus identified two putative pseudogenes for per3 (ENSAMEG00000030567 from giant panda & ENSCJAG00000000288 from White-tufted-ear marmoset), that were removed from further analysis.

Lastly, we removed six species that are not available at TimeTree5 (Sudhir Kumar et al. 2022) and four non-vertebrate species, which additionally four do not have geographic range data on Map of Life/IUCN (last accessed, August 15^th^ 2022). A total of 133 vertebrate species were used for subsequent analyses, species lists and *Per* gene numbers are available in **Table S1**. Amino acid alignments were performed using MAFFT v7 (Katoh et al. 2002). For tree construction, the neighbor-joining distance-based method was applied (Saitou & Nei 1987), focusing on conserved sites across 363 amino acids. The nucleotide substitution model employed was JTT (Jones et al. 1992). Bootstrap analysis was conducted to assess the robustness of the tree, with the number of resampling events set to 100.

### Evolutionary history of gene gain and loss of *Period* genes in vertebrates

We employed a phylogenetic framework that incorporated the species of interest, selected from the TimeTree5 dataset (Sudhir Kumar et al. 2022) (**Supplementary file S2**) accessible via www.timetree.org. The comprehensive tree is the product of a compilation of published molecular trees, the estimation of divergence times, and the construction of a supertree using the hierarchical average linkage method (HAL) of clade pairs; methodological details are available (Hedges et al. 2015). This tree was used to infer the evolution of each *Per* gene in vertebrates. We reconstruct ancestral *Per* states calculating for each gene state transitions based on equal rates (ER), all-rates-different (AR), and symmetrical (SYM) using the hidden-rate-model (Beaulieu et al. 2013) and one category for each level of the trait. The best model was selected according to the Akaike Information Criterion and plotted into the tree, all analyses were performed using ape v5.6 (Paradis & Schliep 2019) and phytools v1.5.1 (Revell 2012) packages. We estimated the phylogenetic signal for each *Per* gene based on Pagel’s lambda λ parameter and the best rate transition for each gene. To examine how *Per* genes changed through time in vertebrates, we used a disparity-through-time (DTT) approach, this analysis quantifies the disparity for the whole clade and each node in the phylogeny (subclade). The relative disparity is calculated by dividing each subclade disparity value by the overall disparity of the clade, whereas the mean relative disparity is calculated for each subclade present at each divergence point. Values close to zero suggest that the variation is partitioned among the distinct subclades in the tree (slowdown in trait diversification), while values near one indicate that a major proportion of the total variation is contained by the subclades in the tree (rapid trait diversification) (Harmon et al. 2003), we performed these analyses in geiger v2.0.10 package (Pennell et al. 2014).

### Association between environmental and ecological factors and Period gene numbers

#### Selection of environmental variables

Besides internal mechanisms, circadian rhythm oscillations in vertebrates are driven by abiotic environmental changes (Zheng et al. 2021). Some of the main environmental factors include the light-dark cycle as the primary factor (Junko et al. 2019), the temperature (Vera et al. 2023), food availability (Vinod Kumar et al. 2022) and oxygen consumption (Adamovich et al. 2022) among others. Based on this information, we chose a set of terrestrial and marine world raster layers as likely factors to predict the evolution of *Per* genes. For terrestrial environments, six climatic layers at 10 arc min (∼340 km²) were obtained from Worldclim v2.1 (Fick & Hijmans 2017): altitude, annual temperature range, annual precipitation seasonality, annual mean solar radiation, annual mean wind speed, and annual mean water vapor pressure. Additionally, we incorporate the Net Primary Productivity of the MOD17A3 v55 layers at 0.1 degrees downloaded from NASA for 2000-2016 (Running et al.), mean value was calculated for each year, and finally, the mean value for the whole period. For marine environments, we selected bathymetry and annual mean temperature range downloaded from MARSPEC at 10 arc min (Sbrocco & Barber 2013), and annual wind speed downloaded at 5 arc min from Global Marine Environment Datasets v2.0 (Basher et al. 2018).

All raster layers were scaled to 10 arc min. The geographic range species maps were obtained from the International Union for Conservation of Nature (IUCN. 2022. The IUCN Red List of Threatened Species v2022-2. https://www.iucnredlist.org. Accessed on October 2022) and Map of Life (Jetz et al. 2012) for extracting raster values and mean values calculated for each species using raster v3.6 and exactextractr v0.9.1 (Baston) in R. The mean latitude was calculated for each species based on the geographic range maps.

#### Selection of ecological variables

Circadian rhythms, shaped by cyclical environmental changes, drive activity patterns in organisms, influencing both physiological and behavioral responses. These activities may be influenced by other factors, including complex ecological factors (Caravaggi et al. 2018). To explore the potential correlation between ecological traits and the evolution of *Per* genes, we accessed ecological trait data for key taxonomic groups from publicly available databases. For fishes, migration, body mass, vision, and swimming mode were downloaded from (Froese et al. 2010) through rfishbase v4.0.0 R package (Boettiger et al. 2012). For amphibians, body mass, habitat, and migration were downloaded from AmphiBio (Oliveira et al. 2017). For birds, body mass, habitat, and migration were downloaded from AVONET (Tobias et al. 2022). For mammals, activity cycle, body mass, and habitat breadth were downloaded from PanTHERIA (Jones et al. 2009).

#### Comparative statistical analyses

To investigate the nature of the evolutionary association between predictors and *Per* gene number we used a Phylogenetic Generalized Least-Squares analysis (PGLS; (Symonds & Blomberg 2014). Analyses were performed for five taxonomic groups/datasets “Mammals” “Amphibians/Reptiles/birds”, “All fishes”, “Continental fishes”, and “Marine fishes”. A violation of the assumptions about the residuals or model instability can severely affect the conclusions drawn, and hence, it is of crucial importance that these are thoroughly checked and an assessment is made about how much the models can be trusted (Mundry 2014). Continuous variables were log10 transformed, for altitude/bathymetry predictor with negative values were squared before log-transformed. The distribution of all transformed variables was checked and tested for collinearity between predictors using the Variance Inflation Factors (VIF) with car package v3.1 (Rasco 2020) before running all models. Variables with no normality and a VIF major of 5 were discarded from the final model.

PGLS models were individually computed for each Per gene using the final predictors, and a phylogenetic signal was determined through Pagel’s lambda (λ), estimated by maximum likelihood (Pagel 1997), and implemented using caper package v1.0.1 (<https://CRAN.R-project.org/package=caper>). Diagnostic plots of residuals, including outlier analysis and Q-Q plots, were thoroughly examined to ensure that assumptions were not violated. A comprehensive overview of the species and the *Per* number, environmental, and ecological data analyzed is provided in **Table S3**.

### Gene-phenotype association datasets

Reported phenotypes associated with *PER1*, *PER2,* and *PER3* in humans at GWAS Atlas (Watanabe et al. 2019) under PheWAS. Traits with a -log_10_ p-value of more than nine in at least one of the *PER* genes were considered to be statistically significant and included in the analysis. Traits in the *Per* deficient mouse were retrieved from the Mouse Genome Database (MGD), 2021 (Blake et al. 2020). Traits in the *per-*deficient zebrafish were retrieved from the Zebrafish Information Network 2022 (zfin.org) (Bradford et al. 2022).

### Transcriptome datasets

The human adult transcriptome data set was retrieved from GTEx portal v8 (last accessed: January 18^th^ 2023) (GTEx Consortium et al. 2017). The mouse transcriptome dataset was retrieved from The mouse Gene Expression Database (GXD) (Baldarelli et al. 2021) Mouse embryo dataset: E-ERAD-401, Mouse adult datasets: E-MTAB-2801 and E-MTAB-6798. Frog (*Xenopus laevis*) transcriptome dataset was retrieved from Xenbase (Fortriede et al. 2020). For adult datasets, we selected organs with available gene expression data in all three species for comparison purposes (muscle, ovary, testis, kidney, liver, spleen, heart, lung, and brain or cerebellum). We used Spearman’s rank correlation coefficient test to investigate the correlation between gene expression patterns between paralogs. For the developmental transcriptome analysis, we used expression data sets from African clawed frog (*Xenopus laevis)* Xenbase (Fortriede et al. 2020), medaka (*Oryzias latipes*) at Medaka Omics Data Portal (Li et al. 2020), and the rhythmicity of gene data sets in Atlantic salmon (*Salmon salar*), *Optic tectum*, gill, and *saccus vasculosus*. Data is retrieved from supplementary table 8 of (West et al. 2020), except for one gene with no expression. For these analyses, we investigated the expression divergence among genes following genome duplication events. Duplicated genes, especially after whole genome duplications, need a rigorous and often species-specific approach to gene annotation. Therefore, for the target species in this analysis that have experienced duplications, we leveraged species-specific datasets for gene subtype annotations where available.

### Molecular evolution analysis

Intensification or relaxation of selection was estimated using RELAX (Wertheim et al. 2015) implemented in HyPhy v2.5 (Kosakovsky Pond et al. 2019). This software compares a set of “test” branches to a set of “reference” branches and measures intensification (k > 1) or relaxation (k < 1) of selection in the test group compared to the reference group. The salmonid group (Atlantic salmon, brown trout) *per3* sequences were used as a test group and compared to sequences of the outgroup (Acanthomorphata) as references.

We explored if there are any sites under diversifying selection on *Per1*, *Per2*, and *Per3* vertebrate genes independently, with MEME (Mixed Effects Model of Evolution) (Murrell et al. 2012) on Datamonkey v2.0 (Weaver et al. 2018). We analyzed each *Per* gene independently as *Per1*, *Per2*, and *Per3* paralogs are too diverged to align and thus expected to reduce statistical accuracy if analyzed simultaneously. MEME estimates a site-wise synonymous and a two-category mixture of non-synonymous rates and uses a likelihood ratio test to determine the signal from episodic diversification, a combination of strength of selection and the proportion of the lineages affected.

### Transcription binding sites analysis

We explored the conservation of E-box elements, expanding previously reported investigations. As general Blast search is not fine-tuned for searching such motif sequences with short query sequences with very rapid turnovers, we used Jbrowse (Buels et al. 2016; Diesh et al. 2022) -based SalmoBase (Samy et al. 2017) (https://salmobase.org/, last accessed, January 5^th^ 2023) to investigate the promoter sequences, allowing ambiguous spacer sequences around the *per* genes in Atlantic salmon. We searched for the three mammalian sequences reported in Nakahata et al. (Nakahata et al. 2008), allowing the six-base spacer sequences for against 100K upstream and downstream within each *Per* gene region.

*per1* caggtcCACGTGcgcccgTGTGTGtgacac *per2*

cgcggtCACGTTttccacTATGTGacagcg *per3*

gaccggCACGCCgcgagcCTCGAGactgcg

We also searched for the predicted CLOCK-controlled cis-elements with E-boxes in figure 5B in Paquet et al (Paquet et al. 2008):

Medaka cgtTCACGTgga-tccccatGTTACA

Zebratish cggTCACCTgtt-tctccacATGCTG

By using Ensembl Comparative Region Comparison Analysis using the 65 fish EPO-Extended chain.

We also conducted an unbiased search using the MEME v5.5.5 (Multiple Expectation maximizations for Motif Elicitation) *de novo* motif discovery results using the 1000 base sequences upstream (the script is available as a Supplementary Material, **file S1**) of each *Per* gene on MEME Suites (Bailey et al. 2015). We used the following parameters for the three *Per* genes:

meme PER1_5prime.fa -dna -oc. -nostatus -time 14400 -mod zoops -nmotifs 10 -minw 18 -maxw 22 -objfun classic -revcomp -markov_order 0.

Our analysis aimed to identify enriched sequences that could potentially serve as cis-regulatory elements. While MEME identified 10 sequences with the lowest p-values, we observed that some of the sequences were merely repetitive sequences, which are likely false positives and unlikely to be cis-regulatory mechanisms. Then, we only considered the reported sequence with the top bits score for each gene.

## Data Availability

We used publicly available datasets, which is described in the Materials and Methods section. Scripts and Supplementary result files are available at FigShare (doi: 10.6084/m9.figshare.25924420).

## Acknowledgments

We express our gratitude to Drs. Shona Wood, Nicola Barson, and Siri Fjellheim for the discussion and comments on the early stage of this study.

## Data Availability

All data used in this work are publicly available, and the accession information is described in the Materials and Methods section.

## Supplementary Materials

**Table S1. Master table of investigated species.**

**Table S2. Excluded species because of “N” in the *Period* genes.**

**Table S3. Variables in continental species.**

**Table S4. Variables in marine species.**

**Table S5. Coefficient of variation of gene numbers in gene families**

**Table S6. MEME selection detection results for *Period* genes.**

**Table S7. The length of each *Period* gene**

**Table S8. NCBI Conserved Domain search results Table S9. MEME motif discovery results**

**Figure S1 RELAX result of salmonid *per3* gene.**

**Figure S2 The distribution of coefficient of gene copy number variation of each gene family across taxa**

**Figure S3 Brain *PER* gene expression in humans.**

**Figure S4 Diversifying selection signature in Period genes in vertebrates.**

**File S1. An R script for gene sequence curation and visualization and fetching upstream sequences**

**File S2. A Timetree-based species tree.**

**File S3. The phylogenetic tree of PER proteins.**

