## Supplementary figures for "Functional and regulatory diversification of *Period* genes responsible for circadian rhythm in vertebrates"

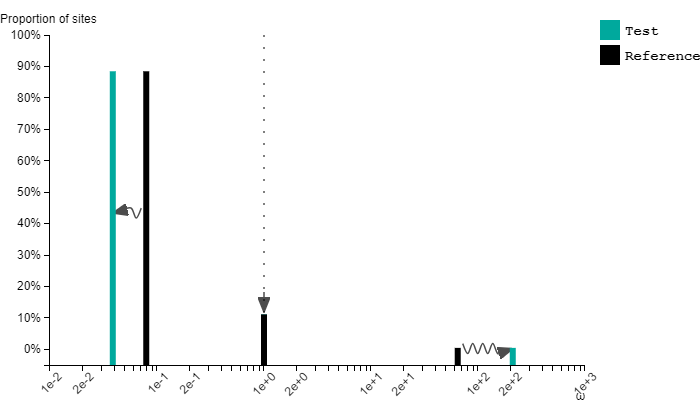

**Supplementary figure S1:** Signature of relaxed selection on per3 sequence in salmon.
Each bar represents the proportion of sites (y-axis) corresponding to the ω values (x-axis) indicated for both the test (green) and reference (black) groups. The arrows indicate the changes in ω values estimated by the model of the software. Figure retrieved from Datamonkey (https://www.datamonkey.org/).

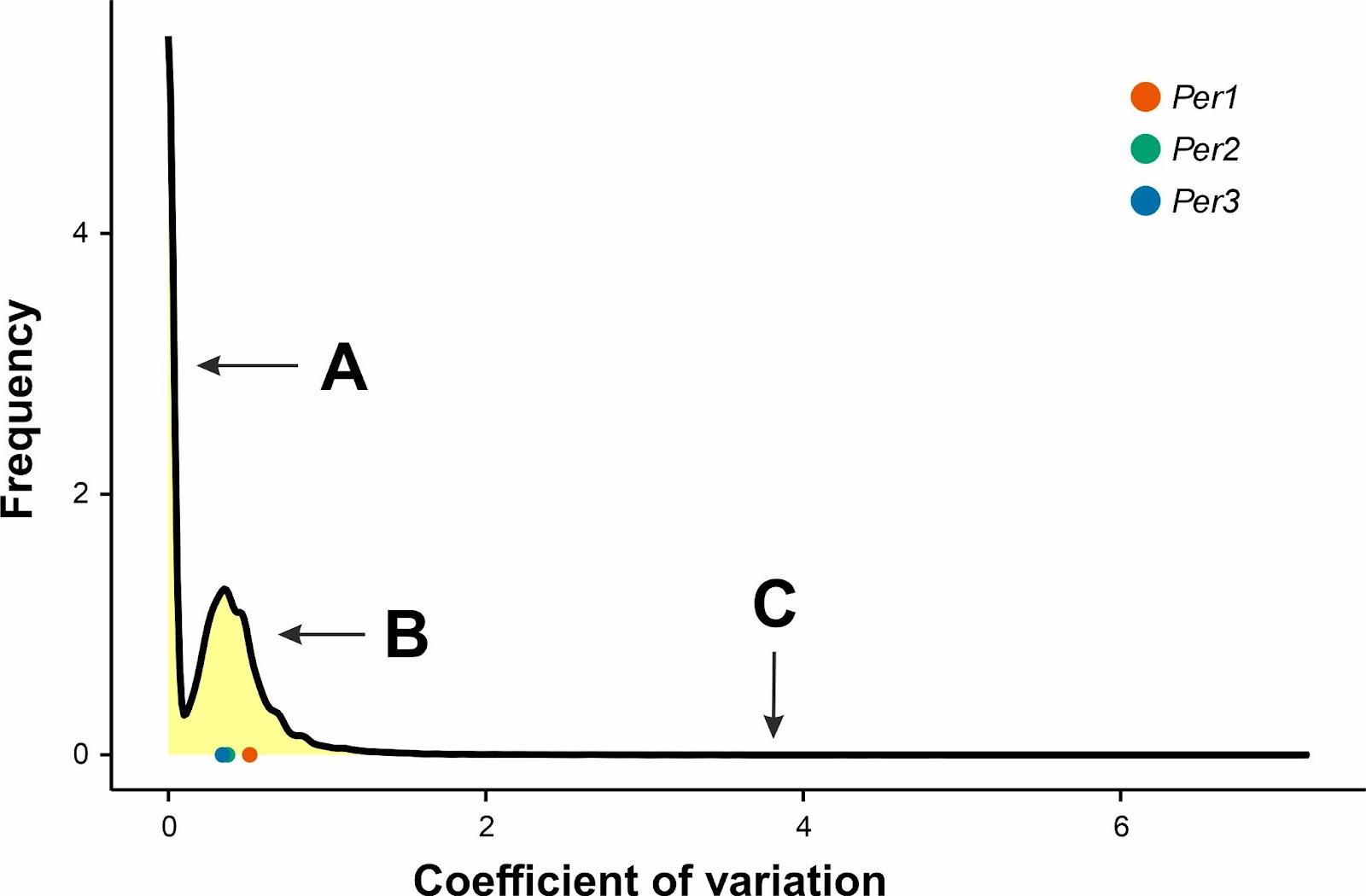

**Supplementary figure 2:** The distribution of coefficient of gene copy number variation of each gene family across taxa. We found three gene groups with very low variation (A), medium variation (B) and high variation (C). per genes fell in the medium variation cluster

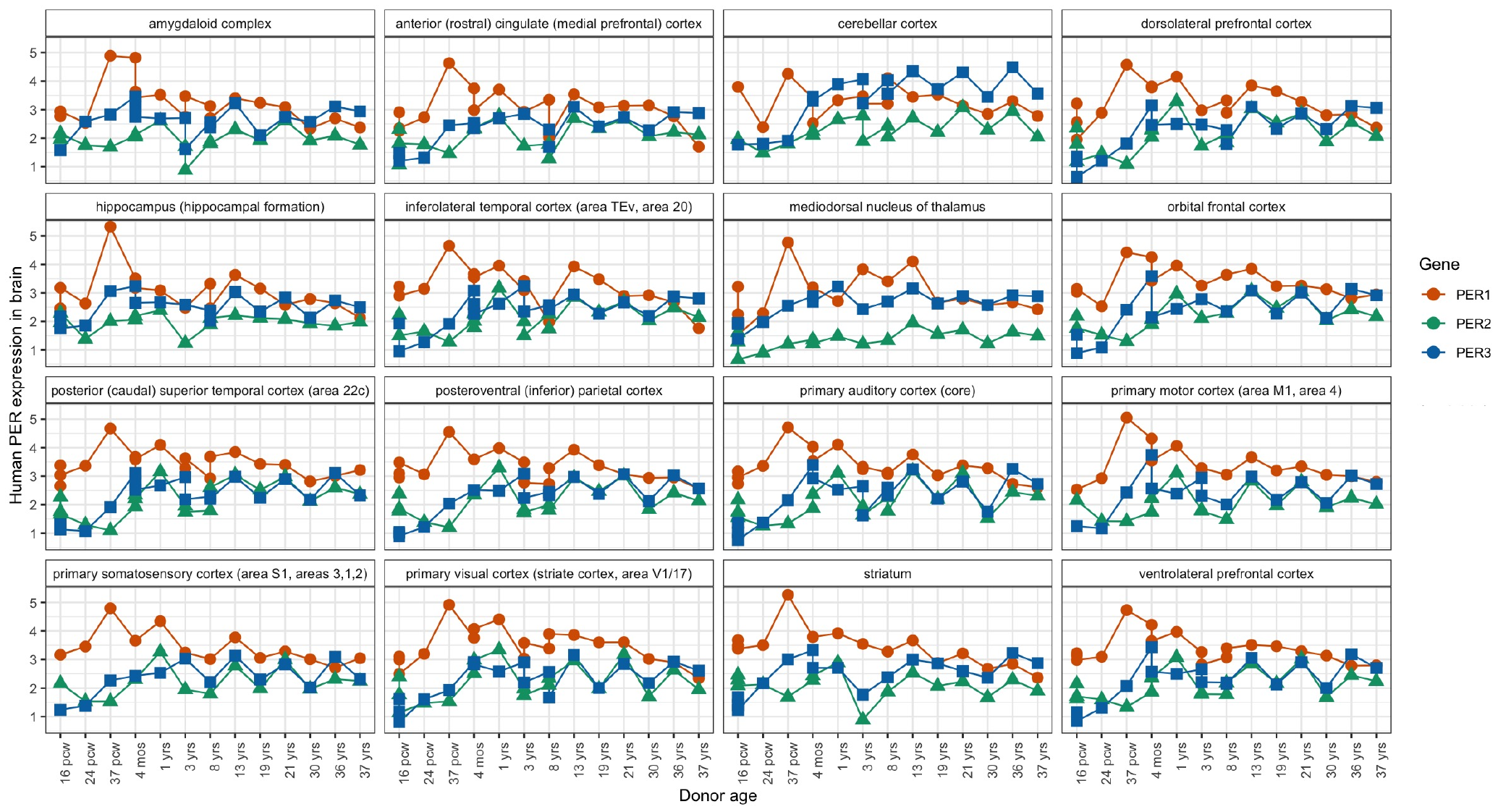

**Supplementary figure S3** Brain *PER* gene expression during human development, retrieved from the data on Brainspan (https://www.brainspan.org/).

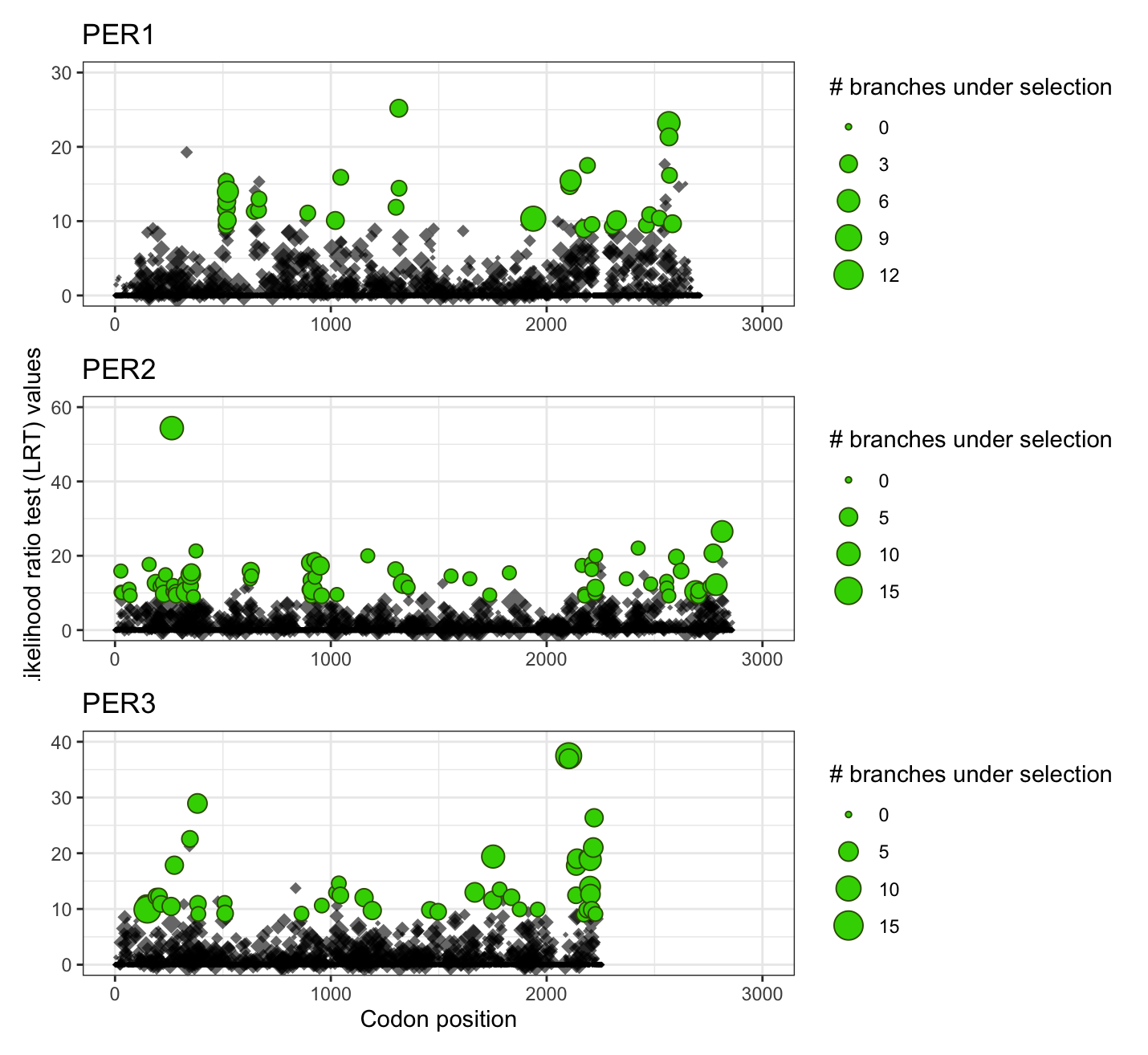

**Supplementary figure S4.** Diversifying selection signature in Period genes in vertebrates detected by MEME (Mixed Effects Model of Evolution) (77) on Datamonkey version 2.0 (78)

Codons with p-value < 0.01 and branches under selection > 1 are highlighted with green dots.
